# Understanding the Influence of Context on Real-World Walking Energetics

**DOI:** 10.1101/2023.06.12.544651

**Authors:** Loubna Baroudi, Kira Barton, Stephen M. Cain, K. Alex Shorter

## Abstract

Speeds that minimize energetic cost during steady-state walking have been observed during lab-based investigations of walking biomechanics and energetics. However, in real-world scenarios, humans walk in various contexts that elicit different walking strategies, which may not always prioritize minimizing energetic cost. To investigate whether individuals tend to select energetically optimal speeds in real-world situations and how contextual factors influence gait, we conducted a study combining data from lab and real-world experiments. Walking kinematics and context were measured during daily life over a week (N = 17) using wearable sensors and a mobile phone. To determine context, we utilized self-reported activity logs, GPS data, and follow-up exit interviews. Additionally, we estimated energetic cost using respirometry over a range of gait speeds in the lab. Cost of transport during these trials was used to identify an energetically optimal walking speed range for each participant. The proportion of real world steady-state stride speeds within this range was identified for all data and for each context. We found that participants walked significantly faster than what was energetically optimal, and the proportion of steady-state strides within the energetically optimal speed range was dependent on the context. On average, 45.6% of all steady-state stride speeds were energetically optimal across all contexts for all participants. These results suggest that while energetic cost is a factor considered by humans when selecting gait speed in daily life, it is not the sole determining factor. Context contributes to the observed variability in movement parameters both within and between individuals.

## INTRODUCTION

Walking is an integral part of daily life, enabling movement in the environment and contributing to overall health and well-being. Studies investigating the relationship between walking kinematics and energetic cost conducted in the lab have demonstrated that humans tend to select movement patterns that minimize cost of transport (Clark 2015). Further, studies in controlled environments have demonstrated that key gait parameters (e.g. step width, step-height, swing limb trajectory, arm motion) are self-selected to minimize energetic cost (Abram et al. 2019; Minetti and Alexander 1997; Zarrugh et al. 1974; Wong et al. 2019; Selinger et al. 2015). However, work has also demonstrated that gait observed in laboratory settings differs from unconstrained movement in the real world (Dal et al. 2010; Lee et al. 2022; Hutchinson et al. 2019; Foucher et al. 2010).

Many of the advantages of in-lab gait analyses, such as increased control over participant behavior, also introduce biases and can make measurements unrealistic. First, although it is favorable for many experiments to study steady-state walking and reduce movement variability, human movement during daily life has been observed to consist of many short bouts or longer walks with punctuated stops from, for example, standing idle waiting to cross a street (Baroudi et al. 2022; Orendurff et al. 2008). Differences have also been observed in walking kinematics in different environments (inside vs outside walking), depending on behavioral context (purposeful vs recreational walks) (Slade et al. 2022; Kim et al. 2020; Wang and Adamczyk 2019). Second, the direct observation of human walking offers greater control during an experiment. However, researchers have shown that individuals alter their performance when observed compared to their natural behavior (a phenomenon known as the “white-coat effect”) (Warmerdam et al. 2020; Hillel et al. 2019).

Compact wearable sensing technologies can be used to measure kinematics and physiology during daily life to investigate walking biomechanics. Small low-power sensors, like accelerometers, have been embedded in shoe soles and belts, and are also commonly found in phones and watches (Benson et al. 2018; Motl et al. 2012). Sensing systems designed around accelerometers can make continuous measurements for weeks (Wu et al. 2022; Baroudi et al. 2022). Inertial measurement units (IMUs) combine accelerometers with gyroscopes and magnetometers to measure additional kinematics. IMUs require more power than an accelerometer alone, limiting recording duration to day scale. But, these sensor data can be used to estimate parameters like foot orientation and speed, enabling position estimates of the foot during gait. The accuracy of the step-to-step position estimates has been shown to be comparable to measurements made using motion capture (L. Ojeda and Borenstein 2007; Rebula et al. 2013; Potter et al. 2019). Using wearables to persistently monitor an individual enables the investigation of real-world movement patterns; however, behavioral and environmental context can also be used to explain the variability observed in real-world movement patterns.

Contextual information can have many forms, from the location of a walk, to its purpose. Prior work has identified a dependency between terrain type and metabolic cost (Gast et al. 2019; Kowalsky et al. 2019). Researchers have leveraged GPS data, as location is closely associated with someone’s activity (Kim et al. 2020; Wang and Adamczyk 2019). Gast, et al. demonstrated how humans might prefer a slower speed than the energetically optimal when walking on difficult terrains to increase stability. However, these experiments were focused on a specific contextual aspect over a short period. To gather contextual information over longer periods, technologies like cameras can offer a direct observation of the environment someone is moving in, but they are highly invasive and the analysis of the images can be burdensome. These issues impact the feasibility of using camera data for long-term monitoring. Alternatively, researchers have asked participants to keep a manual log of their daily activities to gain additional information on context (Cleland et al. 2014; Chang et al. 2017).

The objective of this work is to investigate the relationship between cost of transport, real-world walking speed, and context in young able-bodied adults. Specifically, we will quantify whether the participants used self-selected walking speeds that minimizes their cost of transport in the real world, where walking is highly variable and context-dependent. We hypothesize that, in the real world: (1) Individuals will use on average a walking speed that minimizes their cost of transport, and (2) The use of a walking speed that minimizes cost of transport is independent of context. To test these hypotheses, we combined in-lab modeling to establish the subject-specific relationships between cost of transport and walking speed and long-term data collection to capture real-world walking speed and contextual factors. The results of this study provide insight into the energy expenses of walking in the real world and how contextual information can be effectively used to understand human locomotion. It also highlights the necessity to turn towards ecologically relevant experimental designs for the study of human movement, health, and overall behavior.

## MATERIALS AND METHODS

### Experimental Protocol

#### Recruitment

Data and results from 17 subjects recruited from a healthy population are presented in this work (11 females, 6 males, 26*yo* ± 6*yo*, 168*cm* ± 10*cm*, 65*kg* ± 9*kg*). Participants were excluded if they had health issues that would limit their ability to walk or breath, such as severe cardiac disease or asthma. Participants signed an electronic consent form prior to the first visit, and this study was approved by the Institutional Review Board at the University of Michigan.

#### Real World Data Collection — First Visit

Movement and activity data were first collected over the course of a week from the subjects during their daily life. During an initial visit to the lab, participants were given an accelerometer (activPAL™ [PAL Technologies Ltd., Glasgow, UK]) and an inertial measurement unit (IMU) (Opal, APDM, [Portland, OR, USA]) to wear during the 7 days of monitoring (see Table 1). The activPAL was chosen for its small size and ability to log data continuously for the week-long trial. Additionally, activity classification algorithms developed by PAL Technologies were combined with custom algorithms to isolate walking (Wu et al. 2022; Ryan et al. 2006). Wear time for each sensor can be found in Appendix A.

**Table 1.** Sensor specifications. We extracted walking using the activPAL and derived stride parameters using the IMU.

|  | activPAL | Opal IMU |
| --- | --- | --- |
| Measurement | 3-axis acceleration $\pm 4g$ | 3-axis acceleration $\pm 4g$<br>3-axis angular velocity $\pm 200 \text{ deg} \cdot \text{s}^{-1}$<br>3-axis magnetometer $\pm 8 \text{ Gauss}$ |
| Placement | Thigh | Above shoe on foot dorsum |
| Attachment | Tape | Fabric pouch secured with laces |
| Battery life | 14+ days | 8+ hours |
| Sample rate | 20Hz | 100Hz |
| Size | (23.5 × 43 × 5 mm) | (55 × 40.2 × 12.5 mm) |
| Weight | 9g | 25g |

During the same visit, the Ethica app (Ethica Data [Toronto, Canada]) was installed on participants’ phone. The app logged GPS data from the phone throughout the week. We asked participants to keep the app open in their phone background and to allow the app to access their location. As GPS data can be sensitive, participants were able to turn off the app for periods of time at their discretion (Figure 1 - (A)).

**Fig. 1.**
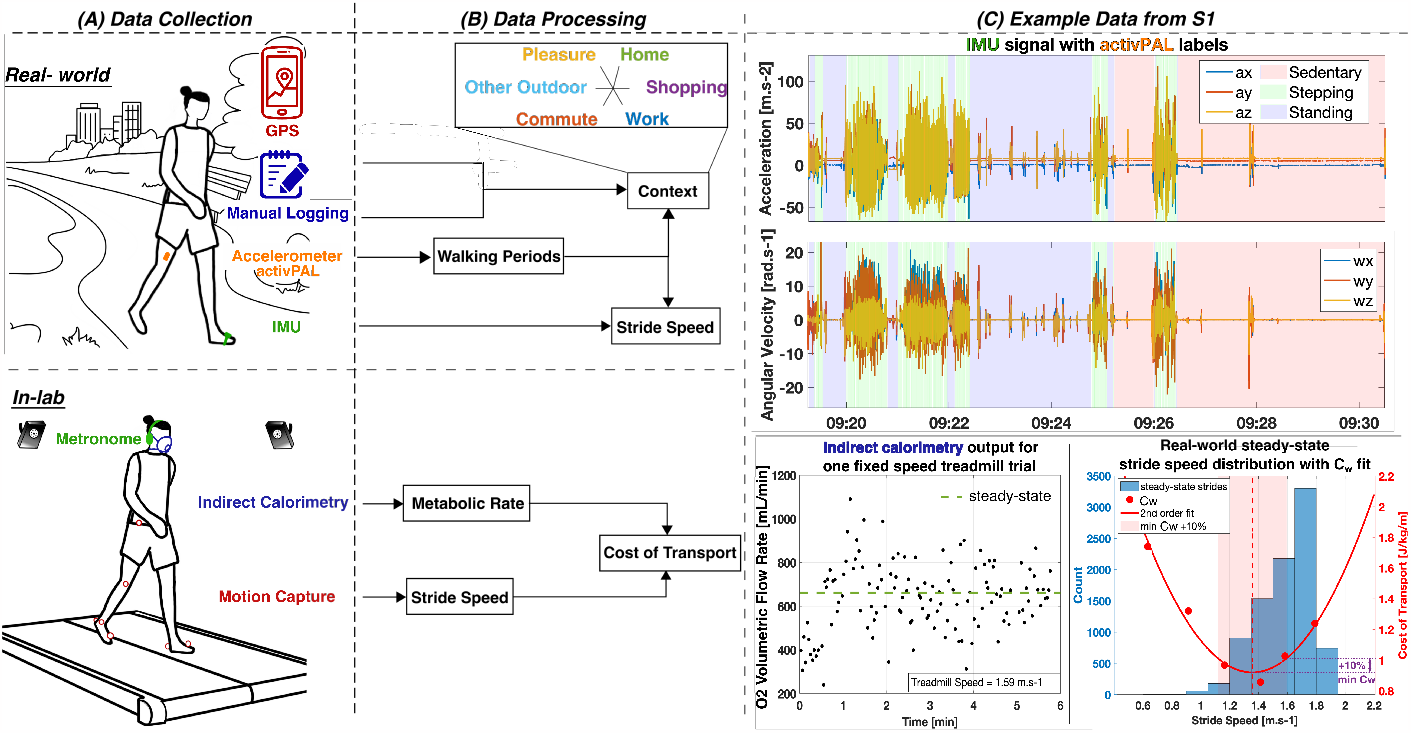
Data Collection and Processing. (A) Data collection combined a week of data collection in the real world and an in-lab treadmill walking task. (B) Data processing methods were used to combine the different sensor streams to obtain stride speed profiles in the real world with associated contexts, and a map of cost of transport vs. stride speed in the lab. (C) The activPAL labels were used to isolate walking periods in the IMU signal in the real world. From these walking periods, we obtained steady-state walking speed profiles. Subsequently, we collected indirect calorimetry data at a range of speed on the treadmill. Then, we built subject-specific models of cost of transport vs. speed to identify the energetically optimal speed range for each individual. The energetically optimal speed range is defined as the minimum cost of transport, plus 10% of that minimum value (shaded region). Finally, we observed the overlap between that range and real-world speeds.

Participants were instructed to go on a daily walk of at least 10 minutes. We specifically prompted participants to take a “pleasure walk” or “stroll”, described as “a walk for the purpose of walking, with no particular purpose nor destination”. This prompt was used to increase context diversity in the data, and ensure that there would be long walking bouts for analysis.

To facilitate labeling and classification, the participants kept an activity log detailing when they were walking, the purpose of the walk, and where they were (e.g., home, work, etc.). A log template and a detailed explanation about how to fill out the log were provided to the participants, along with a set of instructions that included information on how to charge and wear the sensors. Participants could also opt-in to receive daily text reminders to charge and wear the sensors, and to take their daily walk.

#### In-Lab Data Collection — Second Visit

##### Exit Interview

Following the week of real-world data collection, the participants returned the sensors and scheduled the in-lab visit. At the start of the second visit, an exit interview was conducted with the participant to clarify information from the log with the activPAL and GPS data. During the interview, participants were asked to provide supplemental information if the logs were incomplete (e.g. a walk was logged but the purpose is missing). The participants were also asked to recall the context of recorded activities that were not logged (ex: a walk is classified from the sensor data but absent from the log). This interview ensured that the contextual information for the real-world walking data was complete and accurate.

##### Treadmill Walking and Energetics Measurements

An experimental investigation of energetic cost during a range of walking speeds was then conducted with each participant. Sixteen reflective markers were first placed on the participants’ lower body to measure walking kinematics using a 26-camera Vicon Motion Capture system (Vicon, [Oxford, UK]) collecting data at 100 Hz. Markers were placed on the following locations on both right and left leg: Anterior superior iliac, posterior superior iliac, thigh, knee, tibia, toe, ankle, heel. A COSMED K5 system (COSMED, [Rome, Italy]) was used to measure metabolic cost. This system uses a mask worn around the subject’s mouth and nose, and a small backpack that carries the portable measurement unit with the battery. Oxygen and carbon dioxide volumetric flow rate were measured breath-by-breath. To prevent measurement biases, participants were asked to refrain from eating, drinking caffeine, and exercising 5 hours prior to this second visit. Once the participants felt comfortable, they were instructed to perform a 5-min quiet standing to obtain resting metabolic rate. Walking trials were conducted on an instrumented split-belt treadmill (Bertec, [Columbus, OH, USA]) (Figure 1 - (A)). Ground reaction force data were sampled at 1,000Hz from both belts.

Participants were given time to familiarize themselves with the split-belt treadmill before the experiment was conducted. During the experiment the participants were asked to walk on the treadmill at 6 different speeds for 6 minutes each. Data from the week-long data collection was used to determine the range of speeds used for this task. We separated equally in 6 speeds the interval between the maximum and minimum speed observed in the real world. Often, the fastest speed originally chosen had to be reduced to accommodate for the treadmill discomfort that participants felt. The order of the speeds was randomized and a 2-min seated resting period was allowed between each 6-min trial.

We wanted to ensure that participants maintained the walking strategy they used in their daily life on the treadmill to obtain a cost of transport as ecologically valid as possible. However, walking on a treadmill can lead to a decrease of stride length for a given stride speed to increase stability compared to overground (L. V. Ojeda et al. 2015). Thus, we enforced the relationship we found between stride length and stride speed from the real-world walking data for the treadmill trial, by leveraging the relationship between stride length and stride speed. The stride length *l* an individual chooses when walking at a certain stride speed *v* can be predicted for each individual using the following relationship:

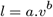

where *a* and *b* are subject-specific model parameters (Grieve and Gear 1966; Kuo 2001). First, we identified the parameters *a*_*rw*_ and *b*_*rw*_ for the real-world model fit for each subject from the data collected prior. Then, we constrained both stride speed and stride length on the treadmill, using the real-world subject-specific models. Stride speed was imposed by the treadmill, but stride length is a difficult parameter to control. Instead, we leveraged the relationship between stride length, stride speed, and stride frequency *f* :

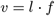

to constrain stride frequency instead. As such, for a given treadmill speed, participants were given a frequency to follow that corresponded to:

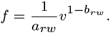

A headset played the calculated metronome frequency and participants were instructed to synchronize their steps to each beat as best as they could. We made sure the participant was able to comfortably match the given stride frequencies on the treadmill, specifically for the highest stride speed. Then, we built the subject-specific models of each participant on the treadmill and found a new set of parameters *a*_*tread*_ and *b*_*tread*_. We evaluated whether this method successfully replicated participants’ real-world walking strategies on the treadmill by comparing the parameters *a*_*rw*_ and *b*_*rw*_ against *a*_*tread*_ and *b*_*tread*_ (Appendix B).

### Data Processing

#### Grouping Real-world Movement Data into Walking Periods

We grouped the real-world walking data into walking periods following the method outlined in Baroudi et al. (Baroudi et al. 2022) (Figure 1 - (B)). Briefly, we used the proprietary algorithm from the activPAL, which identifies stepping (Wu et al. 2022; Ryan et al. 2006). Then, we grouped consecutive stepping bouts into stepping periods if the individual was standing (based on the activPAL classification) for less than 1 minute in between two stepping bouts. The objective of this grouping is to capture the discontinuous nature of real-world walking. For instance, if an individual stops at a pedestrian light before crossing, the walk before and after that stop were grouped together in a stepping period. Lastly, the activPAL algorithm does not distinguish between running and walking within stepping. Thus, we developed a classification algorithm to extract walking periods from all stepping periods.

#### Walking Speed Estimation for a Walking Period

Once we extracted walking periods for each participant, we used the foot-worn IMU data to estimate stride speed (Figure 1 - (B)). Heel strike events in a stride create high peaks in angular velocity that allow for stride detection. The x-axis signal, around which the foot flexes, was smoothed using a locally weighted scatterplot smoothing (LOWESS) method. A peak detection method was applied to isolate strides. Walking periods with less than 5 strides were discarded to reduce errors in gait parameter estimation. To estimate stride speed, we used the zero-velocity update (ZUPT) algorithm (L. Ojeda and Borenstein 2007; Potter et al. 2019; Rebula et al. 2013). This algorithm leverages the assumption that the velocity of the foot on the ground is close to zero when a human is walking to integrate the acceleration while correcting for the IMU drift. The velocity and position of the foot are subsequently obtained. We followed the implementation formulated by Rebula et al. using the same hardware. From the foot position, we extracted stride length and we calculated stride speed by dividing stride length by stride time.

#### Identification of Real-world Steady-state Walking

A custom algorithm (detailed in Appendix C) was used to identify steady-state strides within a walking bout (Figure 2). Briefly, we considered a stride to be at steady-state when the individual was walking straight (e.g. not turning), with a constant velocity (e.g., not accelerating nor decelerating). We started by creating rolling windows of 3 strides in a walking bout with a 2-stride overlap. The second stride of each window was classified as steady-state or non-steady-state using the foot-worn IMU data. The first and last strides of a bout were discarded. Then, for each window, we created a vector with the three stride speeds and stride lengths calculated using the method outlined in the previous sub-section (Walking Speed Estimation for a Walking Period). From these vectors, we calculated the coefficients of variation (*CV*_*speed*_ and *CV*_*length*_) to measure whether stride length and stride speed varied, indicated a possible acceleration or deceleration. If both of these coefficients were above 7%, we classified the stride as non-steady-state. In addition, we calculated cosine similarity between the direction of each pair of consecutive strides. Cosine similarity measures the similarity between two vectors and is bounded between 0 and 1. If the cosine similarity was under 0.7 (e.g., the stride directions are separated by a 45-deg angle), the stride was classified as non-steady-state. Otherwise, the stride was classified as steady-state. The thresholds for the different parameters were determined by conducting a sensitivity analysis as detailed in Appendix C. We evaluated this algorithm using a controlled experiment.

**Fig. 2.**
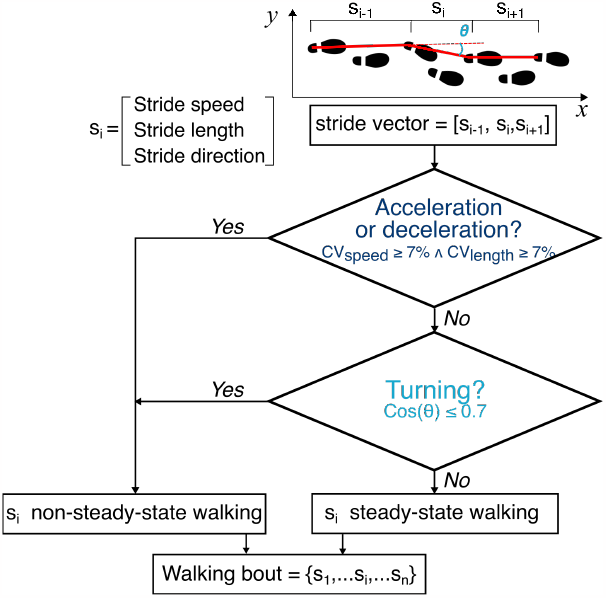
Steady-state identification flowchart. The algorithm and the cut-off choices are detailed in Appendix C. (CV: Coefficient of Variation)

### Real-world Walking Contexts

We manually labeled each walking period using a combination of the manual log given by the participant, the exit interview, and the GPS data (Figure 1 - (B)). For each walking period, we visualized the log entry with a satellite view of the GPS data to isolate the location and purpose of a walk. This process allowed us to isolate repeated locations for a participant, such as their home and workplace. Once all participants’ data were labeled, we isolated six common contexts: *Work, Commute* (e.g., repeated trajectories between home and work), *Pleasure* (e.g., from the pleasure walk that was specifically asked from participants and labeled as such), *Shopping, Home*, and *Other Outdoor* (e.g., any outdoor walk that was not classified as pleasure or commute). Other walks such as church, amusement park, or dog walks were labeled as *Other* as they were rare and often specific to a given participant. The few walking periods (*<* 1%) that had missing labels were also labeled as *Other*.

### In-lab Modelling of Cost of Transport against Stride Speed

We used the force plates from the instrumented treadmill to detect gait events. We discarded strides where the participants did not correctly place their foot on the corresponding belt. The heel marker was used to extract stride length since it moved the least throughout the trials (as opposed to the toe marker for instance that was scrunched during plantar flexion) and never got obstructed from our camera setup. To calculate stride length, we multiplied stride time by the speed of the treadmill belt (assumed constant), and we added the distance between two consecutive foot strikes from the same foot to account for back and forth movement of the participant on the treadmill. Stride speed was then calculated by dividing stride length by stride time.

To obtain metabolic rate at steady state, we used the last 2 minutes of each 6-min trial. The breath-by-breath measurements from the indirect calorimetry system were averaged; then, the Brockway equation was used to estimate whole-body metabolic rate (Brockway 1987). Finally, we subtracted the resting metabolic rate measured during the quiet standing period. To ensure the validity of the derivation, we made sure the respiratory quotient never exceeded 1. We calculated cost of transport by dividing the metabolic rate by stride speed and normalizing by body mass. Lastly, we identified the coefficients of a 2nd order polynomial curve mapping cost of transport to stride speed using the MATLAB curve fitting toolbox. To identify the energetically optimal speed range, we looked at the minimum cost of transport value and added 10%. This resulted in two points on the cost of transport curve, one on the left and one on the right side of the minimum. The associated speeds with these two values of cost of transport were used to define the lower and higher boundary of the energetically optimal speed range. Specifically, the speed corresponding to the point on the left side of the minimum cost of transport represents the lower boundary, while the speed corresponding to the point on the right side represents the higher boundary. By considering the range of speeds between these two points, we can identify an energetically optimal speed range where the cost of transport is minimal. To quantify whether humans used a walking speed within this energetically optimal range, we evaluated the proportion of steady-state strides for each context within the identified range.

For three participants, the measurements of metabolic rate failed for different reasons. One participant’s respiratory quotient was consistently over one. The other did not place the mask properly and had only a limited number of breaths that were registered which did not allow for a sufficiently accurate estimate of metabolic rate. Finally, many participants reported that the mask irritated their eyes and caused nose congestion. One participant could not complete the trial for this reason. We suspect this particular discomfort came from the solution (cidex) used to disinfect the mask between participants.

### Statistical Analysis

A t-test was used to evaluate if participants selected walking speeds that minimize their cost of transport. We tested if the difference between the observed distribution of steady-state stride speeds estimated during daily life and the energetically optimal speeds measured in the lab were significantly different than 0. To test if walking speeds that minimize cost of transport were independent of context, we calculated the percentage of real-world steady-state stride speeds within the energetically optimal speed range for each context and participant. We chose to use a linear mixed model to account for the random effect of participants within each context, given the hierarchical structure of our dataset (context was classified for identified walking periods for each participant) (Field and Wright 2011). We compared different model complexities (with and without random effects) using the Akaike Information Criterion (AIC) (Field and Wright 2011; Akaike 1998) and selected the model that explained the most variance in the data. The tests’ significance were evaluated at the 0.05-level. All statistical analyses were conducted in R.

## RESULTS

### Real-world Walking Behavior

We found 1,731 walking periods across all participants that met our criteria for steady-state (Figure 2 and Appendix C), with a total of 304,309 strides (Table 2). The majority (64%) of walking periods were under 5 minutes, with 15% under 1 minute (Figure 3). The context *Pleasure* contained the most strides (steady-state and non-steady-state) across participants and for almost all individual participants, although S4 and S9 did not take any pleasure walks. The context *Commute* also gathered around 7% of all strides even if only 8 participants had walks in this context (only 45 walking periods total). Gait observed during *Work* contained 25% of the total identified walking periods, but the average duration of these walks were the smallest (2.7min) and represented a little less than 5% of the total identified strides. It is important to note that the proportion of strides within each context was conditioned by the wear of the different sensors. As such, contexts like *Home* may be underrepresented because participants likely remove their shoes (and consequently the foot-worn IMU) when home.

**Table 2.** Walking periods characterization. There are variations between contexts in the number of strides and walking periods, as well as the proportion of steady-state within a walking period. Contexts with longer walking periods tend to have less walking periods, but more strides and steady-state strides.

|  | # Strides | # Walking Periods | # Steady-state strides | Avg steady-state strides within walking period [%] | Avg Walking period duration [min] |
| --- | --- | --- | --- | --- | --- |
| Other Outdoor | 90,292 | 228 | 72,210 | 71.9 | 11.4 |
| Pleasure | 95,506 | 115 | 82,520 | 82.4 | 20.9 |
| Commute | 21,082 | 45 | 18,181 | 85.9 | 12.6 |
| Work | 14,173 | 431 | 7,524 | 48.4 | 2.7 |
| Shopping | 12,752 | 143 | 8,176 | 50.9 | 5.1 |
| Home | 15,338 | 344 | 5,681 | 30.7 | 4.3 |
| All Walks | 304,309 | 1,731 | 230,515 | 51.7 | 6.5 |

**Fig. 3.**
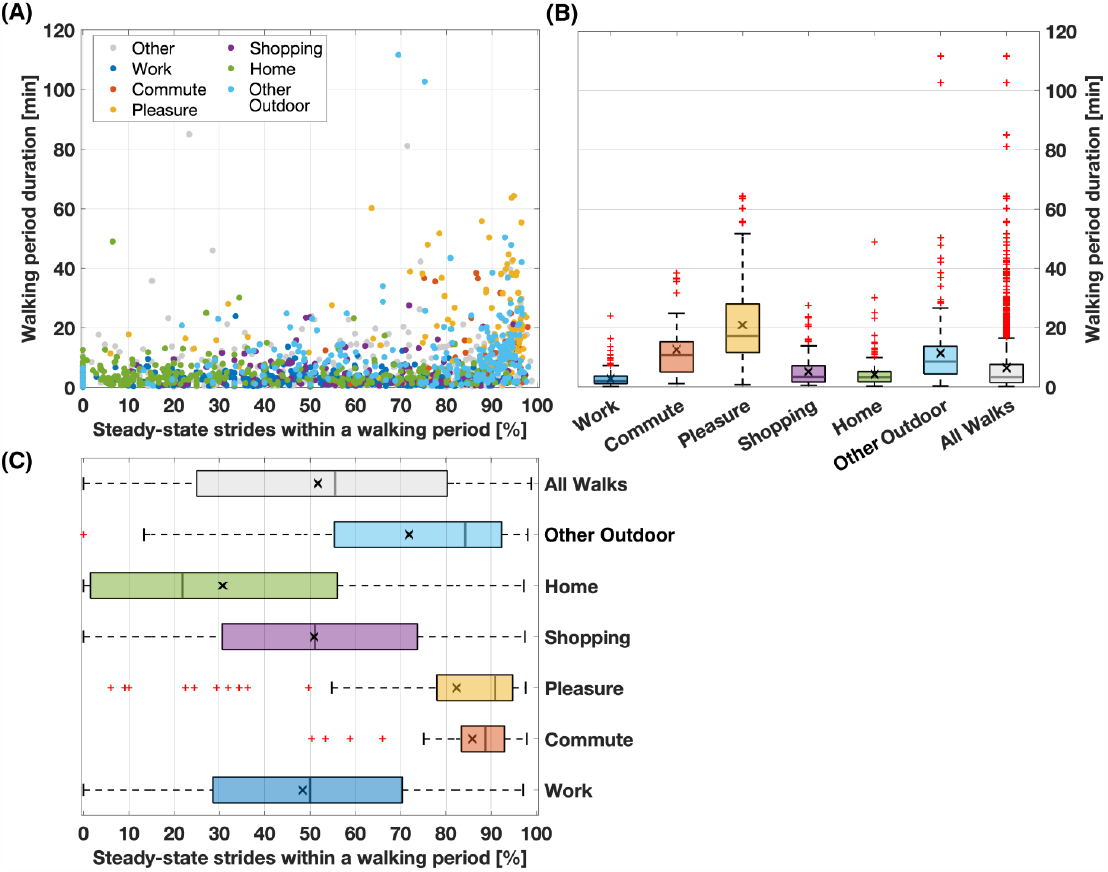
Walking periods duration and their proportion of steady-state strides. Relationship between walking period duration and percentage of steady-state strides within a walking period — (A) Each dot corresponds to a walking period. The different colors are for each context. (B) We binned all walking periods by their bout duration and looked at the percentages of steady-state strides within a walking period for each bin. (C) We binned all walking periods by their percentage of steady-state strides and looked at the walking periods durations for each bin.

### In-lab Cost of Transport Models

We mapped cost of transport to walking speed using data collected during the in-lab treadmill walking task (Figure 1 - C). The models had an average *R*^2^ of 0.78 and RMSE of 0.20*J/kg/m*. Energetically optimal speeds that minimized cost of transport varied between individuals (Table 4 and Figure 5), ranging from 0.82*m* · *s*^*−*1^ for S8 to 1.37*m* · *s*^*−*1^ for S4 with an average minimum cost of 1.83*J/kg/m*.

### Steady-state Walking and Context

Overall, an average of 51.7% of the identified strides were classified as steady state, but the percentage of steady-state strides varied by context. Around 30% of the strides identified during walking at home were steady state, while 86% of the strides during a commute were steady state (Figure 3). We observed high variability in the proportion of steady-state strides during walking in indoor contexts (e.g., *Home, Shopping*, and *Work*) (Figure 3). On the other hand, these particular contexts had a low variability in walking period duration with averages between 2.7 minutes for *Work* and 5.1 minutes for *Shopping*. Figure 4 presents a summary of the identified steady-state strides based on both context and energetically optimal stride speeds. We found that contexts containing longer duration walking periods, such as *Commute* or *Pleasure*, also contained a higher percentage of steady-state strides (32.7% and 31.6% of all steady-state strides respectively).

**Fig. 4.**
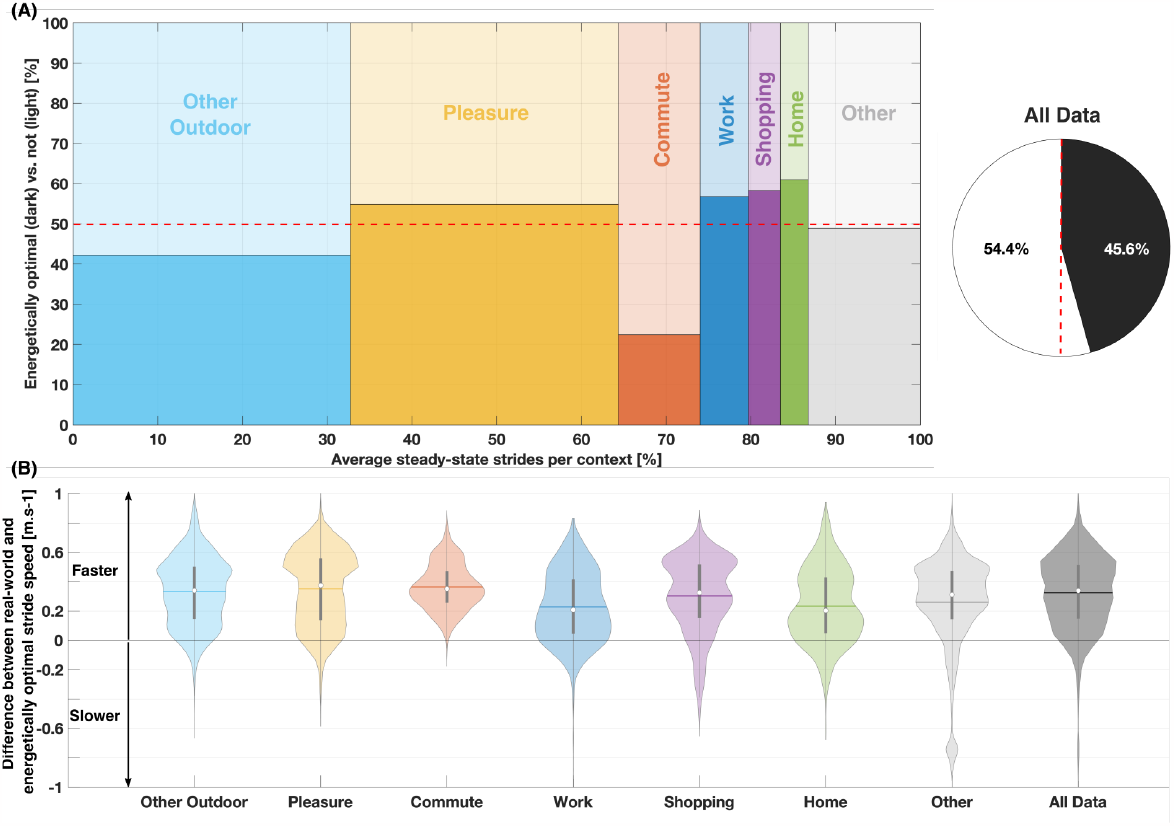
Energetic optimality for steady-state strides in each context. (A) Proportion of steady-state strides for each context for all participants with the associated proportion of strides within the energetically optimal speed range. The darker parts of each bar represents the proportion of strides with speeds within the energetically optimal range. For instance, we read that the context Pleasure represents approximately 35% of the data and that a little more than half of the strides have a stride speed within the energetically optimal speed range. (B) Distribution of stride speed subtracted by the subject-specific speed at the minimum cost of transport for all subjects in each context. Each shaded area represents the shape of the distribution, and the horizontal lines marks the mean. For instance, we read that participants walked on average 0.35*m · s*^*−*1^ faster than their energetically optimal when commuting.

### Cost of Transport Minimization and Walking Speed Variability

Figure 4 — A) shows the proportion of energetically optimal speeds for each contexts and for all data. The distribution of the different between real-world and energetically optimal stride speed for all participants combined is shown in Figure 4 — B). Figure 5 shows this same distribution but for each individual in each context. We tested whether our participants walked in the real world at speeds comparable to the energetically optimal speeds identified in the lab. We found that, across all participants and contexts, participants walked at a stride speed significantly higher than the energetically optimal (*p <* 0.05), with a mean difference of 0.32*m* · *s*^*−*1^ (30% increase). However, a large percentage of the steady state strides were still at energetically optimal speeds. Overall, participants walked within the energetically optimal range for 45.6% ± 33.0% of their strides (Figure 4 and Figure 5). Next, we tested whether there was a dependence between the use of an energetically optimal walking speed and context. We found a significant relationship between the proportion of energetically optimal speeds (e.g., real-world stride speeds within the energetically optimal speed range) and contexts across participants, *SD* = 0.29 (95% CI: 0.19, 0.42). All contexts, apart from *Other Outdoor*, significantly predicted the proportion of strides at energetically optimal speeds. *Commute* was arbitrarily chosen as the baseline for the model. The coefficients indicate that participants were more likely to use energetically optimal speeds in all contexts when compared to *Commute* (Figure 4). All of the coefficients are reported in Table 3. Individuals walked much faster when commuting relative to both the other contexts (*∼* +0.14*m · s*^*−*1^) and to the energetically optimal speed range (*∼* +0.36*m · s*^*−*1^).

**Table 3.** Linear Mixed Model Coefficients

|  | *b* | *SE b* | *p-value* |
| --- | --- | --- | --- |
| *Commute* | 28.1 | 10.6 | 0.010 |
| *Home* | 34.0 | 9.7 | 0.001 |
| *Other Outdoor* | 13.1 | 9.6 | 0.098 |
| *Pleasure* | 29.3 | 9.9 | 0.004 |
| *Shopping* | 27.1 | 10.65 | 0.014 |
| *Work* | 31.3 | 10.0 | 0.003 |

**Table 4.** Subject-specific data. Each participant had different economical speed range, leading to varied proportions of economical walking speeds in the real world for the different contexts. For each context, participants displayed different walking strategies as well.

| Eco speed range |  | Work |  |  | Commute |  |  | Pleasure |  |  |
| --- | --- | --- | --- | --- | --- | --- | --- | --- | --- | --- |
| | | # Walking Periods | # steady-state strides | Stride Speed ($\mu$, $\sigma$) | # Walking Periods | # steady-state strides | Stride Speed ($\mu$, $\sigma$) | # Walking Periods | # steady-state strides | Stride Speed ($\mu$, $\sigma$) |
| S1 | [1.12,1.36,1.59] | 43 | 774 | (1.46,0.16) | 4 | 5027 | (1.70,0.10) | 1 | 532 | (1.34,0.06) |
| S2 | [0.63,0.86,1.10] | 0 | 0 | - | 0 | 0 | - | 2 | 3329 | (1.40,0.07) |
| S3 | [0.74,1.00,1.26] | 37 | 1104 | (1.43,0.20) | 11 | 977 | (1.49,0.12) | 1 | 787 | (1.27,0.07) |
| S4 | [1.11,1.37,1.62] | 0 | 0 | - | 1 | 344 | (1.57,0.06) | 0 | 0 | - |
| S5 | [0.73,1.02,1.31] | 16 | 531 | (1.54,0.13) | 1 | 257 | (1.59,0.10) | 18 | 13506 | (1.67,0.08) |
| S6 | [0.71,1.08,1.44] | 118 | 1275 | (1.09,0.12) | 0 | 0 | - | 10 | 4653 | (1.09,0.13) |
| S7 | [0.97,1.27,1.27] | 13 | 352 | (1.27,0.15) | 0 | 0 | - | 8 | 7784 | (1.30,0.09) |
| S8 | [0.37,0.82,1.28] | 11 | 220 | (1.24,0.12) | 0 | 0 | - | 8 | 9661 | (1.35,0.06) |
| S9 | [0.43,0.94,1.46] | 21 | 326 | (1.36,0.10) | 2 | 450 | (1.60,0.10) | 0 | 0 | - |
| S10 | [0.81,1.09,1.37] | 11 | 173 | (1.48,0.19) | 4 | 2044 | (1.54,0.10) | 6 | 3863 | (1.42,0.11) |
| S11 | [0.84,1.05,1.26] | 9 | 121 | (1.23,0.09) | 3 | 2532 | (1.34,0.08) | 9 | 11013 | (1.30,0.10) |
| S12 | [0.96,1.16,1.37] | 0 | 0 | - | 0 | 0 | - | 9 | 2119 | (1.14,0.09) |
| S13 | [0.84,1.18,1.53] | 44 | 981 | (1.37,0.14) | 0 | 0 | - | 5 | 927 | (1.18,0.10) |
| S14 | [0.66,0.96,1.26] | 0 | 0 | - | 0 | 0 | - | 3 | 2837 | (1.39,0.11) |
| S15 | [0.78,1.07,1.37] | 60 | 715 | (1.27,0.15) | 13 | 5103 | (1.41,0.19) | 4 | 2123 | (1.28,0.12) |
| S16 | [0.56,0.89,1.22] | 0 | 0 | - | 0 | 0 | - | 8 | 3552 | (1.35,0.31) |
| S17 | [0.80,1.08,1.36] | 2 | 1 | (1.63,0) | 0 | 0 | - | 1 | 550 | (1.29,0.08) |

| Eco speed range |  | Shopping |  |  | Home |  |  | Other Outdoor |  |  |
| --- | --- | --- | --- | --- | --- | --- | --- | --- | --- | --- |
| | | # Walking Periods | # steady-state strides | Stride Speed ($\mu$, $\sigma$) | # Walking Periods | # steady-state strides | Stride Speed ($\mu$, $\sigma$) | # Walking Periods | # steady-state strides | Stride Speed ($\mu$, $\sigma$) |
| S1 | [1.12,1.36,1.59] | 9 | 174 | (1.47,0.23) | 9 | 264 | (1.48,0.08) | 21 | 2254 | (1.46,0.19) |
| S2 | [0.63,0.86,1.10] | 31 | 755 | (1.29,0.17) | 11 | 59 | (1.11,0.33) | 10 | 8240 | (1.39,0.15) |
| S3 | [0.74,1.00,1.26] | 13 | 333 | (1.29,0.21) | 11 | 28 | (1.24,1.16) | 9 | 2740 | (1.50,0.07) |
| S4 | [1.11,1.37,1.62] | 0 | 0 | - | 3 | 5 | (1.16,0.03) | 10 | 4286 | (1.53,0.09) |
| S5 | [0.73,1.02,1.31] | 0 | 0 | - | 6 | 22 | (1.55,0.20) | 19 | 3811 | (1.62,0.19) |
| S6 | [0.71,1.08,1.44] | 6 | 588 | (1.23,0.18) | 10 | 592 | (1.18,0.12) | 2 | 3215 | (1.20,0.10) |
| S7 | [0.97,1.27,1.27] | 13 | 316 | (1.14,0.15) | 13 | 212 | (1.28,0.09) | 0 | 0 | - |
| S8 | [0.37,0.82,1.28] | 10 | 1994 | (1.38,0.08) | 32 | 150 | (1.36,0.12) | 14 | 7443 | (1.36,0.11) |
| S9 | [0.43,0.94,1.46] | 0 | 0 | - | 0 | 0 | - | 2 | 515 | (1.49,0.08) |
| S10 | [0.81,1.09,1.37] | 5 | 237 | (1.44,0.14) | 7 | 124 | (1.39,0.09) | 27 | 6157 | (1.52,0.13) |
| S11 | [0.84,1.05,1.26] | 0 | 0 | - | 8 | 22 | (1.23,0.09) | 19 | 10499 | (1.36,0.12) |
| S12 | [0.96,1.16,1.37] | 11 | 360 | (1.12,0.15) | 1 | 9 | (1.20,0.04) | 47 | 10980 | (1.19,0.11) |
| S13 | [0.84,1.18,1.53] | 8 | 497 | (1.16,0.21) | 5 | 32 | (1.40,0.14) | 6 | 1526 | (1.44,0.12) |
| S14 | [0.66,0.96,1.26] | 0 | 0 | - | 82 | 753 | (0.96,0.15) | 0 | 0 | - |
| S15 | [0.78,1.07,1.37] | 0 | 0 | - | 5 | 7 | (0.65,0.14) | 4 | 2444 | (1.42,0.13) |
| S16 | [0.56,0.89,1.22] | 5 | 76 | (1.01,0.16) | 21 | 1333 | (1.33,0.20) | 7 | 1543 | (1.51,0.41) |
| S17 | [0.80,1.08,1.36] | 15 | 1741 | (1.35,0.10) | 26 | 345 | (1.28,0.16) | 6 | 1456 | (1.34,0.09) |

The values and the variability of walking speed changed from one context to another for the same participant (Table 4 and Figure 5). For example, during commutes and pleasure walks, S1 exhibited low variability, but largely different walking speeds with a more energetically optimal walking speed used during pleasure walks (*μ* = 1.70 ± 0.10*m* · *s*^*−*1^ and *μ* = 1.34 ± 0.06*m* · *s*^*−*1^, respectively). On the other hand, when shopping or at work, the variability in the walking speeds was larger with a greater overlap with the energetically optimal speed ranges (*μ* = 1.47 *±* 0.23*m · s*^*−*1^ and *μ* = 1.46 *±* 0.16*m · s*^*−*1^, respectively).

**Fig. 5.**
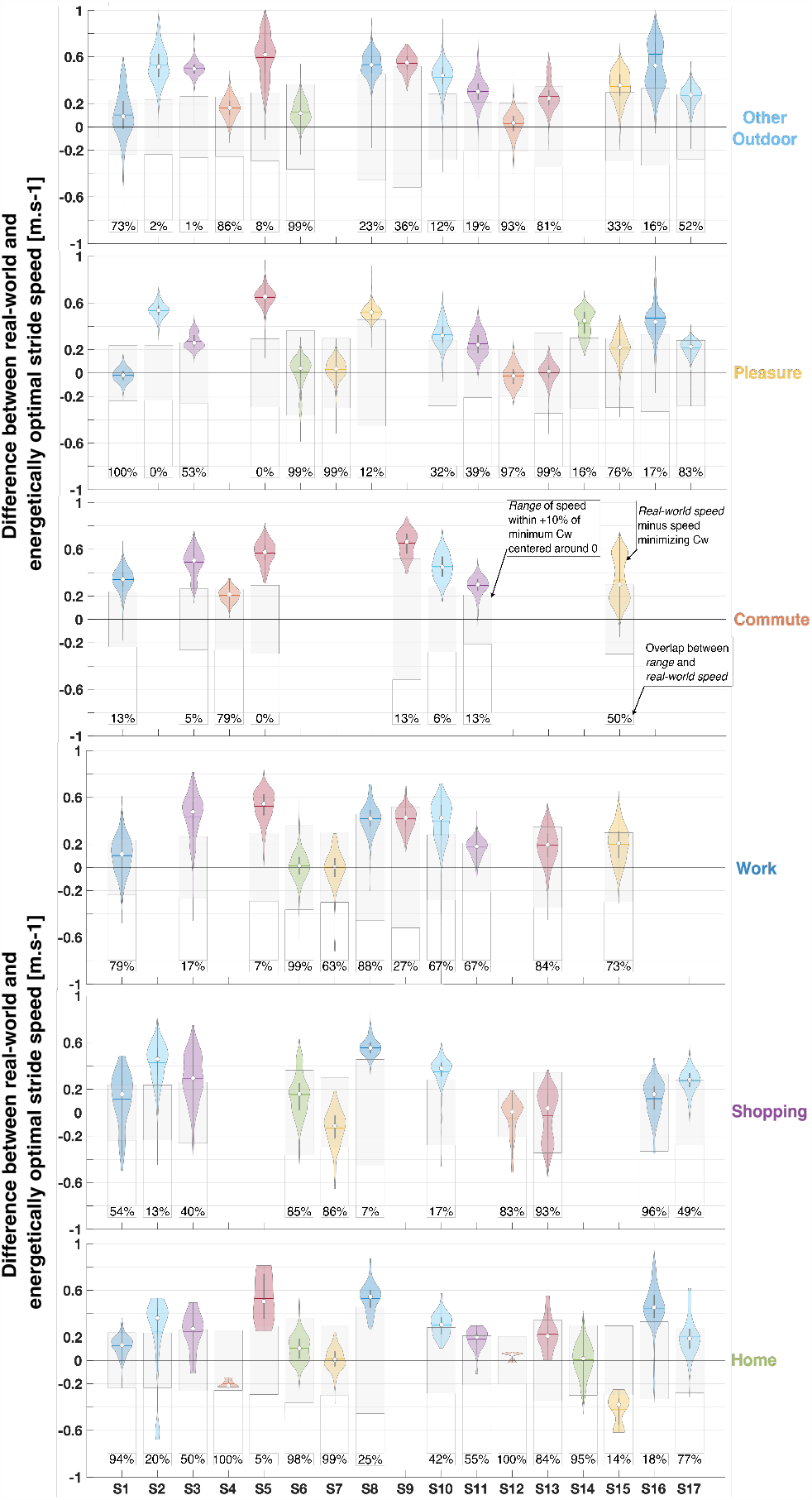
Steady-state stride speed distributions and proportion of energetically optimal strides for each context. Distribution of stride speed subtracted by the speed at the minimum cost of transport for the different contexts. Each shaded area represents the shape of the distribution for a subject, with the horizontal line for the mean. The energetically optimal speed range is centered around 0 and represented by the grey boxes. The percentages represent the proportion of speeds within the energetically optimal speed range. Some individuals had no data for a given context.

## DISCUSSION

Laboratory-based studies have found that individuals tend to use walking speeds that minimize cost of transport. Building on these results, the work presented here investigated walking economy in the real world by testing two hypotheses: (1) on average, individuals will self-select energetically optimal walking speeds and (2) the use of energetically optimal walking speeds is independent of context. We rejected both hypotheses as we found that: (1) participants most often used faster walking speeds than what is energetically optimal and (2) the proportion of walking speeds within the energetically optimal range varied based on context. This suggests that individuals have multiple competing objectives when walking in the real world, including minimizing cost, which influences walking strategy.

### Real-world Walking Economy, Context, and its Applications

Various in-lab studies have observed a convergence of different kinematics parameters towards an energetically optimal movement pattern (Abram et al. 2019; Wong et al. 2019; Selinger et al. 2015). Most of these studies were conducted in the lab, during steadystate walking, where participants walked for bouts shorter than 20 minutes. As such, we expected to see a similar relationship in the real world, particularly during long walks at steady state. First, we found that individuals walked approximately 30% faster in the real world compared to their energetically optimal speed across all contexts. Second, the amount of energetically optimal speeds that an individual used was found to be dependent on context.

We observed that the predominantly indoor contexts (*Shopping, Work, Home*) contained the most energetically optimal strides. In these contexts, physical constraints like halls, doors, rooms, and furniture may affect how individuals are able to move in the environment. While only around 12% of all steady-state strides were observed during these contexts, these constraints (e.g., with many starts/stops, and turns) might have led participants to slow down and walk at a stride speed closer to their energetically optimal (Figure 4 and Figure 3).These contexts also resulted in a high variability in walking speeds, which might have led to a bigger overlap with the energetically optimal speed range (Figure 4 and Table 3). We expected participants to select energetically optimal speeds in the contexts *Other Outdoor, Pleasure*, and *Commute*, as they contained longer walking periods with more steady-state behavior (Figure 3). These three contexts contained around 74% of all data. We found that 55% of the steady-state walking speeds during a *Pleasure* walk were energetically optimal, compared to 42% and 22% for *Other Outdoor* and *Commute* respectively. It is important to note that we specifically asked participants to take “pleasure walks” to elicit a walk without any particular destination or objective other than walking. Therefore, the higher proportion of energetically optimal strides within this context may reflect the results from laboratory-based studies as there were no other competing objectives. In contrast, the other outdoor walks and commuting on foot often have fixed destinations and time constraints (e.g., getting to work on time), which could explain the use of faster walking speeds that have a higher energetic cost. This difference was particularly clear for commuting, where the highest walking speeds were measured (Table 3).

These findings can provide insight into research aiming to develop technological advancements based on human energetics, such as research in lower-body assistive devices. In fact, these fields of research often consider energy minimization as the main optimization objective. However, our results suggest that, depending on the context in which someone is walking, they may not prioritize walking at energetically optimal walking speeds. These walking speeds could still be optimal, but for a different objective than energy minimization. It is possible that, given the many constraints of our real-world environment, individuals cannot necessarily use predominantly energetically optimal behavior. As such, experimental designs should evolve to take into account real-world scenarios in order to ensure the ecological relevance of technological innovations. In Slade et al., a subject-specific optimization of a lower-body exoskeleton was carried out in the real world (Slade et al. 2022). The researchers recreated a realistic real-world scenario where participants walked short bouts in an uncontrolled environment, using “ecologically relevant audio prompts” that elicited variability in the walking speed used by the participants. Such experimental design ensures that the technology (here an exoskeleton) can be used by individuals across different contexts and walking behaviors.

It is difficult to quantify the potential energetic consequences or penalties on the metabolism that someone experiences or perceives when deviating from their energetically optimal walking patterns. Across all individuals, we observe deviations in stride speed from the energetically optimum up to 0.65*m* · *s*^*−*1^ (e.g., 57% more from min cost of transport for S5 in the context *Pleasure*). In Medrano et al., they estimated the “Just Noticeable Difference” in metabolic rate that individuals perceived with the assistance of a lower-body exoskeleton walking on a treadmill (Medrano et al. 2022). They found that participants (N = 10) were able to perceive changes of around 20% ± 5% with an accuracy of 75%. Thus, individuals may be mostly unable to perceive the deviations we observed from the energetically optimal walking speed. Even if participants could perceive the deviation, the energetic penalty could potentially be negligible given that most walks were relatively short in distance and duration (Figure 3).

### Steady-state Walking in the Real World

In the animal kingdom, the best locomotion strategy depends on context and might not always involve steady-state. A prey evading a predator utilize transient states with high acceleration that prioritize maneuverability over energetic consumption (Moore et al. 2017). In this work, a large portion of the observed movement occurred during non-steady-state/transient behavior (48.3% of all strides were classified as non-steady-state), although paticipants did not have to evade predators (as far as we know). The proportion of steady-state strides was dependent on context. As expected, walks in more spatially constrained contexts (ex: at home) contained more non-steady-state behavior (Figure 3) (Glaister et al. 2007). Although fewer strides were identified in these contexts (compared to *Commute* or *Other Outdoor* for instance), they contained more walking periods. In other words, when an individual stood up to walk, they were more often inside, in contexts such as *Work* or *Home* (22.2% and 14.4% of walking periods, respectively). This prevalence of transient behavior highlights the importance of studying all types of human movement that can be observed in the real world, particularly for health and mobility. For example, LA King et al. leveraged the importance of turns and the potential risks associated for people with Parkinson’s Disease when turning (King et al. 2022).

### Variability in Walking Strategy

We observed large variability within the identified steady-state strides between, and within individuals. Between individuals, we noted a large variability in the overlap between the energetically optimal speed range and the real-world speed distribution within the same context (Figure 5). Overall, individuals used walking speeds that were faster than energetically optimal, but there did not seem to be a way to predict the overlap size for a given context for all participants. For instance, in the context *Pleasure*, the overlap ranged from 0% to 100%. This suggests that other variables could potentially explain the variability between individuals, such as fitness level. Second, within individuals, we observed a large variability in the walking speed used for different contexts (Figure 5 and Table 3). For instance, both *Commute* and *Other Outdoor* contained long walking periods with a large proportion of steady-state strides (Figure 3). However, we measured differences in walking speed up to 0.24*m* · *s*^*−*1^. Preferred walking speed is a clinically important parameter used to quantify health and well-being (Fritz and Lusardi 2009; Graham et al. 2008). The variability observed in our data makes it challenging to estimate preferred walking speed in the real world. Speed estimated during a *Pleasure* walk is the least likely to be influenced by context, and may be the most comparable to data collected in the lab. However, future work should investigate whether leveraging the ensemble of the data and deriving a usable range of speeds to characterize preferred walking behavior could be a stronger health indicator. Taken together, these variabilities in walking strategy confirm that researchers should consider designing experiments that allow participants to display the range of behavior they naturally use in the real world. Additionally, the development of subject-specific models to capture an individual’s natural movement profile could lead to more accurate representations of an individual’s health and well-being.

### Limitations and Potential Opportunities

Our experimental design combined in-lab and real-world measurements. We built subject-specific energetically optimal speed ranges during in-lab measurements on the treadmill, and compared these ranges to real-world measurements. However, to mitigate the effect of the differences between the laboratory and the real world, we: 1) enforced the relationship between stride speed and stride length found in the real world on the treadmill (Appendix B), and 2) used a large uncertainty interval of +10% around the minimum cost of transport (which creates a range of speeds that correspond to the minimum of the cost of transport curve) to capture possible differences between the lab and the real world. Additionally, future research should broaden the population diversity and increase the sample size. The methods and insights from this study could potentially be relevant for clinical populations. Finally, we defined context as a combination of location and purpose; however, there are many external contextual factors we did not include, such as terrain type or topography. Future studies should also consider internal contexts, such as an individual’s mood or fatigue, as they may play a role into someone’s movement in the real world.

## CONCLUSION

Researchers now have the ability to measure human movement in the real world. Persistent monitoring is a powerful tool that can create rich datasets and provide a unique understanding of natural human behavior. In this work, we found that humans walk in a variety of contexts and exhibit a wide range of walking strategies and behavior. Consequently, individuals do not necessarily use a walking speed that minimizes their cost of transport in the real world. Overall, we recommend that researchers take into account the possible discrepancies between in-lab and real-world behavior when studying walking biomechanics. For instance, in-lab experiments can be designed to simulate real-world conditions to fully represent an individual’s movement profile and capabilities. We believe the findings of this work can provide insight for improvements in the study of human mobility for health, but also in robotics for the design of lower-body assistive devices.

## APPENDIX A SENSOR WEAR TIME

Participants were instructed to wear the activPAL during the whole week but were allowed to remove it overnight if they wished. As such, the compliance for this sensor was high. We had an average of 6h of non-wear time for the week across all participants. S1 accumulated the most non-wear time over the week (36.4h). They removed the sensor for an entire day and one night for personal reasons. 13 participants had less than 5h of non-wear time over the weak.

The IMU was attached on participants shoe using a pouch secured with their laces. As such, participants were instructed to wear the IMU whenever they went out and were wearing their shoes. On average, participants wore the sensor for 9.2h per day. The average daily wear time ranged from 6.9h for S3, to 13.2h for S12. Many participants were able to recharge the sensor during the day, which explains high daily wear time. Sundays were the days with the least wear time (8.4h). Some participants did not wear the sensor for an entire day, either because they forgot to place the sensor in their shoe, or because they did not wear that specific pair of shoe, or because they did not leave their home on that day.

## APPENDIX B ENFORCING REAL-WORLD WALKING STRATEGY ON THE TREADMILL

As explained in the materials and methods (Experimental Protocol - In-Lab Data Collection — Second Visit - Treadmill Walking and Energetics Measurements), we constrained participants’ stride frequency for a given treadmill speed to enforce the parameters *a* and *b* of the power model:

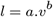

Here, we show the results of the difference in the parameters we found in the real world and those the participants displayed on the treadmill.

We found less than 10% change in parameter *a* apart for S12 and S15. There was more change in parameter *b*, with an average difference of 23%. For 10 participants, the change in parameter *b* was an increase. For 15 participants, the change in parameter *a* was a decrease. This reflects the tendency of participants to lower their stride length for a given stride speed, and particularly for faster speeds. In fact, when *b* increases, stride length decreases for a higher stride speed, and when *a* decreases, stride length decreases for all given stride speeds. Figure B1 shows an example with 2 different participants real-world vs. treadmill power models. S7 adopted a consistently lower stride length for a given stride speed when walking on the treadmill, whereas S12 matched successfully their in-lab walking to the real world.

## APPENDIX C STEADY-STATE WALKING IN THE REAL WORLD

The notion of steady state can be used to describe any type of dynamic process that reaches an unvarying condition. For human movement, steady-state walking is the convergence of specific parameters to a point of stability. The most common characterizations of steady-state walking use either physiological parameters (e.g., heart rate, oxygen consumption) (Parvataneni et al. 2009), mechanical parameters (e.g., stride speed, stride frequency) (Macfarlane and Looney 2008; Strutzenberger et al. 2021; Lindemann et al. 2008), or a combination of both. In the real-world, the large variability of contexts can lead to a range of walking behavior involving a combination of steady-state and non-steady-state. Slowing down, stopping, starting, or turning are all motions that can lead to non-steady-state gait. In our study, we wanted to extract steady-state walking from real-world data using our set of sensors. We used mechanical parameters to characterize and differentiate steady-state and non-steady-state walking.

### The Algorithm

The algorithm (detailed below) was applied to all walking bouts (within walking periods) for a specific subject. We iterated through all strides (N) within a bout and discarded the first and last strides. We started by creating vectors of stride speeds and stride lengths using three consecutive strides (*s*_*i*−1_, *s*_*i*_, and *s*_*i*+1_). The coefficient of variation *CV* was calculated for each of these vectors (*CV*_*length*_ and *CV*_*speed*_). A high coefficient of variation is indicative of large variations from the mean in a given vector. In our case, it would indicate a change of stride speed or stride length within the three selected strides. Then, we calculated the direction of the strides *s*_*i*−1_ and *s*_*i*_ using the foot position derived from the integration of the IMU data as explained in the sub-section (Walking Speed Estimation for a Walking Period). Cosine similarity *cosθ* was calculated, with *θ* being the angle between the direction of *s*_*i*−1_ and *s*_*i*_. Cosine similarity was chosen to evaluate the alignment in the direction of the two consecutive strides. If *cos*(*θ*) is close to 1, it means the vectors are almost aligned. Finally, we compared *CV*_*length*_, *CV*_*speed*_, and *cosθ* against thresholds to classify the stride *s*_*i*_. The choice of thresholds is explained in the next section.

#### Algorithm 1 Steady-state classification of strides

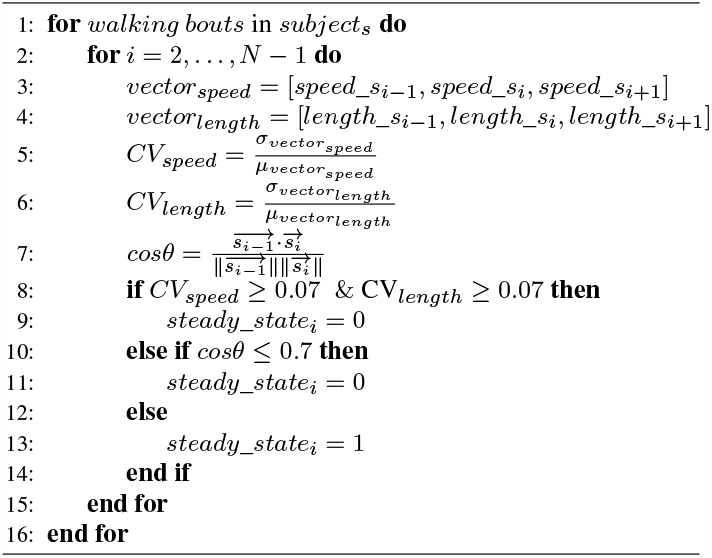

### Sensitivity Analysis for Threshold Selection

To determine the thresholds for the three parameters we selected, we conducted a sensitivity analysis following the leave-one-out method. We varied one parameter while fixing the two others to explore how sensitive our outputs were to each variable. The outputs we monitored were: (1) the distribution of steady-state stride speed for each participant, and (2) the percentage of strides identified as steady state across all participants. We fixed *CV*_*speed*_ and *CV*_*length*_ to 0.07 and 0.07 respectively. Those fixed values were derived from Macfarlane et al. (Macfarlane and Looney 2008). We fixed *cos*(*θ*) to 0.7, which translates to a maximum of 45 degrees between the direction of two consecutive strides.

**Fig. B1.**
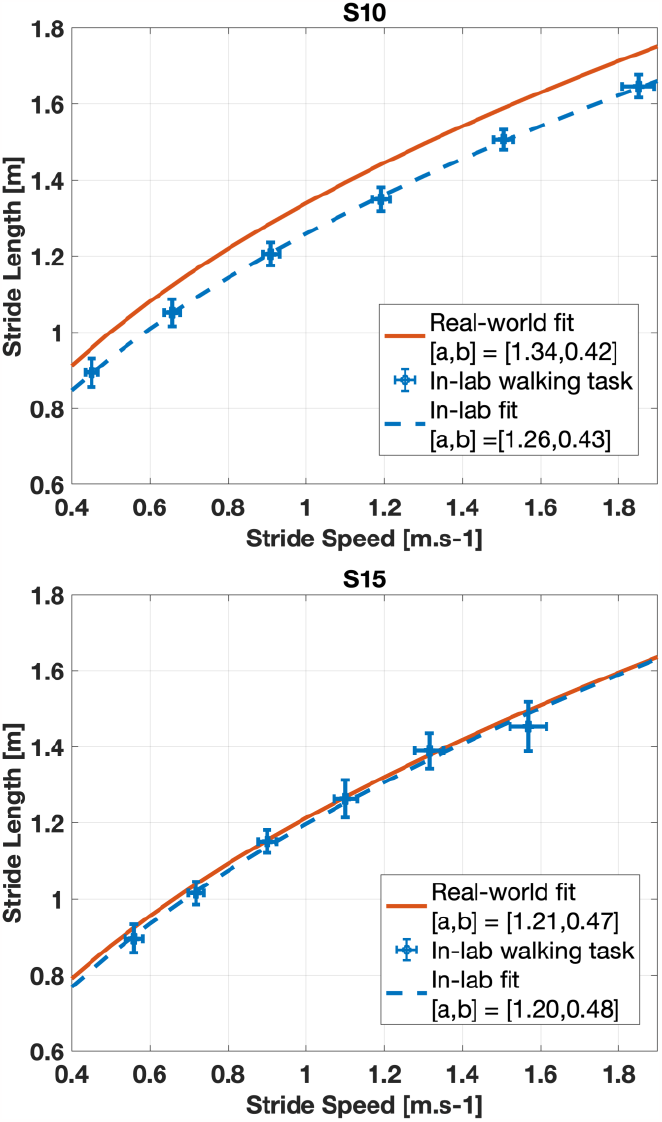
Enforcing Walking Strategy. Power model mapping stride length to stride speed for both the real-world and the in-lab data collection. Each dot represents the average for a given trial on the treadmill and the errorbars represent the standard deviation for both stride speed and stride length. We chose to not show the real-world data points to not crowd the plots.

We did not observe a large sensitivity of the average and standard deviation of steady-state stride speed when varying the different parameters. Thus, we did not use this output to select the thresholds for each parameter.

In Figure C1 - (A) and (B), we observe that the variation of both *CV*_*speed*_ and *CV*_*length*_ influences the percentage of identified steady-state strides following a first-order response trend, with a convergence to a value close to 80% for a representative subject. On the other hand, the variation *cos*(*θ*) did not influence the percentage of identified steady-state strides apart for very large values (e.g., *cos*(*θ*) *>* 0.97) (Figure C1 - (C)). Humans generally vary their stride speed and stride length when they are about to turn. Thus, this result indicates that most turns in the real world are likely captured by a change of *CV*_*speed*_ and/or *CV*_*length*_. We used the results from the sensitivity analysis on this output to determine thresholds for each parameter. Since *cos*(*θ*) did not influence either outputs in a meaningful way, we kept a threshold of 0.7. For *CV*_*speed*_ and *CV*_*length*_, we picked the value at which the percentage of identified steady-state strides crossed the ± 2% boundary around the value this output converged to (Figure C1 – (A) and (B)). This resulted in a choice of 0.07 for both *CV*_*speed*_ and *CV*_*length*_.

### Validity of Thresholds

To confirm the validity of the thresholds we chose, we used a controlled in-lab experiment in which participants walked back and forth on a hallway. We reconstructed the track of each participant and observed the identified steady-state strides using our selected thresholds. Figure C2 shows a representative subject’s track with the associated changes in parameters *CV*_*length*_, *CV*_*speed*_, and *cos*(*θ*). Steps around a turn to walk back in the hallway were identified as non-steady-state. We also notice that the second stride and the second to last stride were classified as non-steady-state. This corresponds to the transition between walking and stopping. As a reminder, the first and last strides were discarded to be able to run the designed algorithm. Each parameter shows how the accelerations and decelerations as well as the turns are captured by the different chosen thresholds (Figure C2 - (B)).

## ACKNOWLEDGEMENTS

The authors thank the Neurobionics Lab and the Locomotor Control Systems Lab of the University of Michigan for sharing their space with us for this study.

## COMPETING INTERESTS

No competing interests declared.

## FUNDING

This research was supported by the Precision Health Initiative at the University of Michigan and the Patricia C. Schroeder Family Fund Award.

**Fig. C1.**
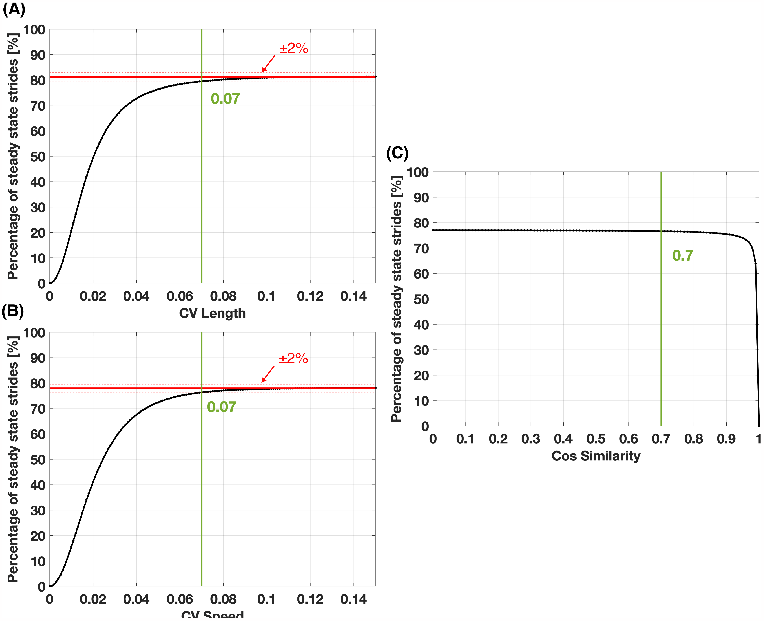
Sensitivity Analysis. Variation of the percentage of steady-state strides based on changes in the parameters *CV*_*length*_ (A), *CV*_*speed*_ (B), and *cos*(*θ*) (B) for a given subject.

**Fig. C2.**
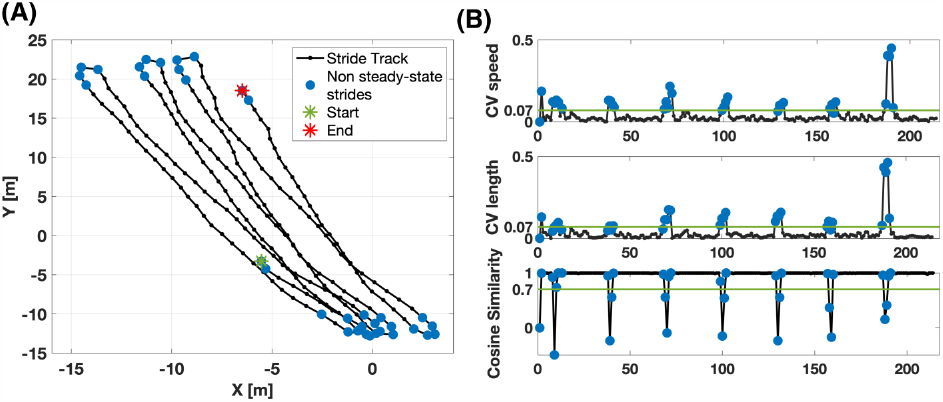
In-lab Walk for Validation. (A) Representative track for S7 during a controlled in-lab walk where participants walked back and forth in a hallway. (B) Parameters variation associated with the track. The green line represent the chosen threshold for each parameter.

## Notes

### Competing Interest Statement

The authors have declared no competing interest.

